# Intergenerational pathogen-induced diapause in *C. elegans* is modulated by *mir-243*

**DOI:** 10.1101/2020.03.31.019349

**Authors:** Carolaing Gabaldon, Marcela Legüe, M. Fernanda Palominos, Lidia Verdugo, Florence Gutzwiller, Andrea Calixto

## Abstract

The interaction and communication between bacteria and their hosts modulate many aspects of animal physiology and behavior. Dauer entry as a response to chronic exposure to pathogenic bacteria in *Caenorhabditis elegans* is an example of a dramatic survival response. This response is dependent on the RNAi machinery, suggesting the involvement of sRNAs as effectors. Interestingly, dauer formation occurs after two generations of interaction with two unrelated moderately pathogenic bacteria. Therefore, we sought to discover the identity of *C. elegans* RNAs involved in pathogen-induced diapause. Using transcriptomics and differential expression analysis of coding and long and small non-coding RNAs, we found that *mir-243-3p* is the only transcript continuously upregulated in animals exposed to both, *P. aeruginosa* or *S. enterica* for two generations. Phenotypic analysis of mutants showed that *mir-243* is required for dauer formation under pathogenesis but not under starvation. Moreover, DAF-16, a master regulator of defensive responses in the animal and required for dauer formation was found to be necessary for *mir-243* expression. This work highlights the role of a small non-coding RNA in the intergenerational defensive response against pathogenic bacteria and inter-kingdom communication.

**Importance:** Persistent infection of the bacterivore nematode *C. elegans* with bacteria such as *P. aeruginosa* and *S. enterica* makes the worm diapause or hibernate. By doing this, the worm closes its mouth avoiding infection. This response takes two generations to be implemented. In this work, we looked for genes expressed upon infection that could mediate the worm diapause triggered by pathogens. We identify *mir-243-3p* as the only transcript commonly upregulated when animals feed on *P. aeruginosa* and *S. enterica* for two consecutive generations. Moreover, we demonstrate that *mir-243-3p* is required for pathogen-induced dauer formation, a new function that has not been previously described for this miRNA. We also find that the transcriptional activators DAF-16, PQM-1 and CRH-2 are necessary for the expression of *mir-243* under pathogenesis. Here we establish a relationship between a small RNA and a developmental change that ensures the survival of a percentage of the progeny.

## Introduction

*C. elegans* has a close evolutionary relationship with bacteria (1) as it has naturally evolved exposed to microbes from the soil that can either be their food source or a threat (2, 3, 4). In laboratory settings *C. elegans* has been fed for decades on the standardized bacteria *E. coli* OP50 and just in recent times our understanding of this nematode-bacteria relationship has evolved from a simple static organism/substrate pair to a dynamic model in which host and microbe’s performance changes throughout their association (5). Moreover, their ability to recognize and defend themselves from potential pathogens has likely been shaped by its continuous encounters with different types of bacteria, and thus when confronted with infectious microbes, *C. elegans* can avoid them by displaying complex behavioral, endocrine and immune responses (6, 7). The worm response is triggered by specific molecules secreted by bacteria such as toxic pigments from *P. aeruginosa* PA14 (8); cyanide from *P. aeruginosa* PAO1 (9); and serrawettin W2, produced by *S. marcescens* (10); but mediated by ASJ and ASI neurons through the activation of the DAF-7/TGF-β pathway (7) in concert with neuro-peptidergic control of innate immunity (11) to finally modify their olfactory preferences (6). When avoidance is not possible and worms are exposed to highly pathogenic bacteria they die within 24 hours (12). In contrast, when confronted with mild pathogenic bacteria for two or more generations, a percentage of the population enters diapause forming the dauer larvae, an alternative stress-resistant larval stage that does not feed, thus is able to properly avoid pathogen infection (13). As pathogen-induced dauer formation (PIDF) depends on the RNA interference (RNAi) machinery (13) we propose that for its initiation PIDF requires the expression of a specific set of small RNAs (sRNAs), and at the long-term, the maintenance of sRNAs expression across more than one developmental cycle. This accumulation of sRNAs could generate molecular footprints that will predispose the upcoming generations of worms to enter diapause thus ensuring the survival of a percentage of the total population. In this way, the sustained but dynamical communication between host and pathogen enables the worm’s development and reproduction over consecutive generations (13).

The molecules signaling diapause entry in the second generation after pathogen exposure are unknown. This work focuses on the discovery of sRNAs involved in the response of *C. elegans* to chronic pathogenic infection that leads to defensive dauer formation. Here we show that *mir-243-3p* is overexpressed in animals exposed to two unrelated pathogens and is needed to mount intergenerational pathogen-induced diapause formation. We also show that transcription factors DAF-16, PQM-1, and CRH-2 are required for the expression of the mature form of *mir-243.* Furthermore, PQM-1 and CRH-2 are also needed for dauer formation under pathogenesis. This work reveals an intergenerational role for *mir-243* in the defense against pathogens and highlights the importance of small RNAs as mediators of long-term survival strategies.

## Results

### Global gene expression changes in intergenerational chronic exposure to pathogens

Developmental and behavioral plasticity likely emerges from broad, complex gene expression changes at different molecular levels. To reveal RNA profiles underlying the intergenerational diapause entry (13) we performed a transcriptomic analysis of two generations of synchronized *C. elegans* in the L2 stage. Animals were grown on the two diapause-inducing bacteria *P. aeruginosa* PAO1 and *S*. *enterica* serovar Typhimurium MST1; and on *E. coli* OP50, which does not trigger dauer formation. We aimed at finding transcriptomic changes elicited by both pathogens in the two generations that could explain PIDF. Sequencing was performed on both mRNA and small RNA libraries generated by separate methods (see Materials and Methods). In this first result, we will address polyA+ RNAs while sRNAs will be addressed in the next section. To detect mRNAs and other polyadenylated transcripts, mRNA libraries were polyA-selected (polyA+). We performed differential gene expression analysis of animals feeding on pathogenic bacteria, using non-pathogenic *E. coli* OP50 as a reference. We considered differentially expressed (DE) those genes with a log2 fold change of >1, padj <0.05 by DeSeq and p <0.05 by EdgeR (Dataset 1; Table S1 and 2). Differentially expressed polyA+ RNAs included coding and non-coding transcripts (**Figure 1A-D**). Among non-coding transcripts, we found piRNAs, 7k ncRNA, pseudogenes, tRNAs, lincRNAs, asRNAs, snoRNA, snRNA, and rRNAs (**Fig. 1E** and **F**). Surprisingly, transcriptional changes in coding and non-coding polyA+ RNAs were much larger in abundance and diversity in the first (F1) than in the second generation (F2) of animals fed with either pathogen (**Fig. 1A-F**).

**Fig. 1.**
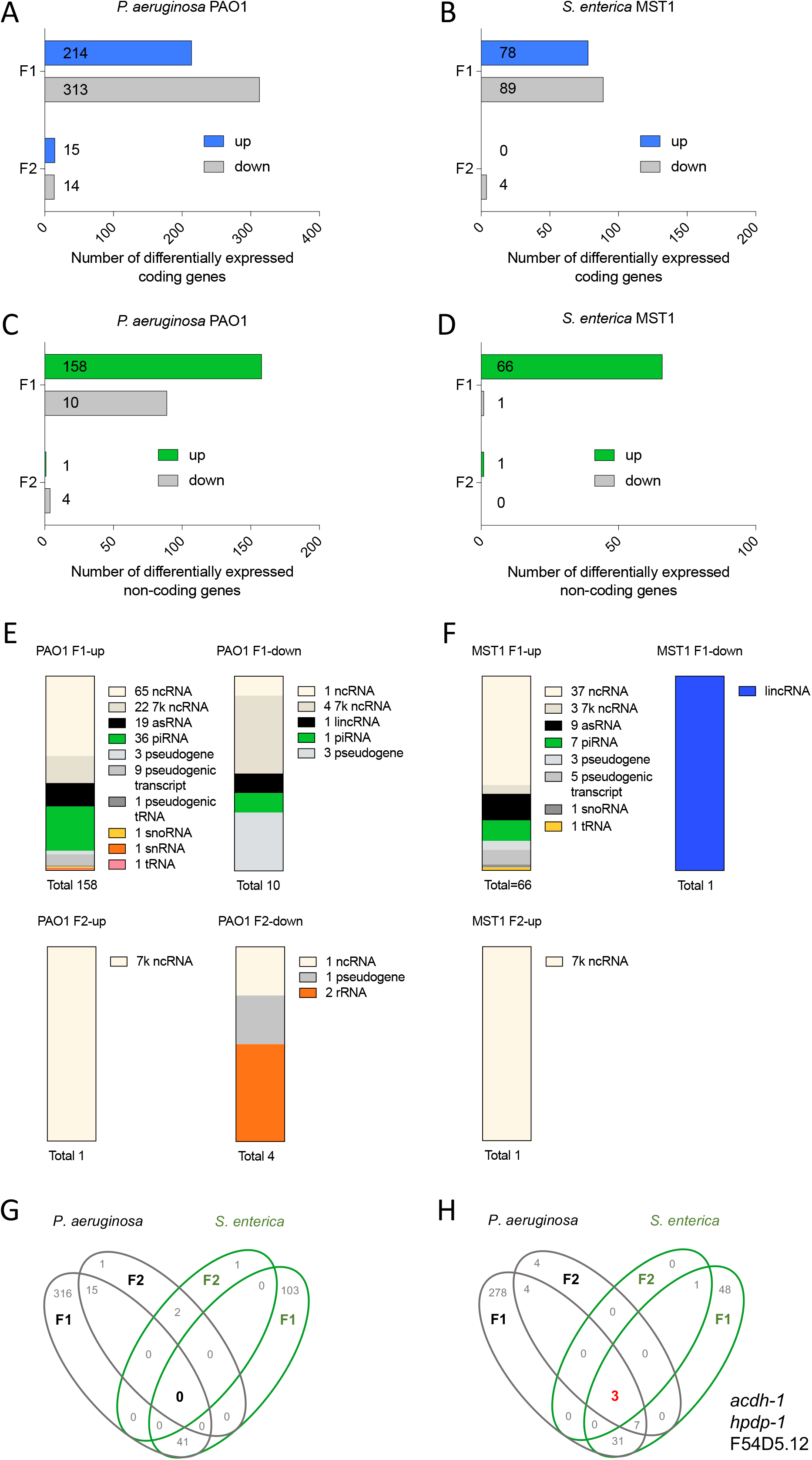
Global analysis of differential mRNA gene expression of an intergenerational infection paradigm. **A-D**. Number of polyA-RNA coding (A, B) and non-coding (C, D) genes differentially expressed on *P. aeruginosa* PAO1 (A, C) and S. *enterica* serovar Typhimurium MST1 (B, D) in two generations. **E-F.** Type and abundance of non-coding RNAs on *P. aeruginosa* PAO1 and *S*. *enterica* MST1 in two generations. **G-H**. Venn diagram representation of shared and unique genes up (G) and down (H) regulated in each generation and on each pathogen.

PolyA+ RNAs were overexpressed or repressed in a generation and/or pathogen-specific manner (**Table S3**) and did not share upregulated coding or non-coding polyA+ sequences that were common to both pathogens and both generations (**Fig. 1G**). Notwithstanding, we found coincidences for the repression of three genes: *acdh-1, hphd-1,* and F54D5.12 (**Fig. 1H**), in both pathogens and across the two generations. All coding genes differentially expressed in the F2 were also up or downregulated in the F1, with no new genes of this kind turned on or off selectively in the F2 (**Table S3**).

To distinguish between changes caused by food switch from *E. coli* to other bacteria from changes induced by long-lasting pathogenic exposure, we compared the expression profiles of animals feeding on *P. aeruginosa* and *S*. *enterica* in the F1, to those reported for *Comamonas aquatica*, a nourishing food for *C. elegans* (14). Upregulated coding genes were only present between pathogens (“pathogen-exclusive” in **Fig. 2A**), but repressed genes were found in worms exposed to the three bacteria (**Fig. 2B**). These 16 downregulated genes were all enriched in gene ontology (GO) terms related to metabolic processes. This reveals a common “food switch factor” caused by changing diet from *E. coli* to other food sources despite their pathogenic potential. We then compared the expression changes produced in the second generation (F2) of worms exposed to *P. aeruginosa* and *S. enterica* to those produced by worm’s first encounter with *C. aquatica* (14). Surprisingly, *acdh-1, hphd-1,* and F54D5.12 (**Fig. 1H**) remained downregulated in both pathogens and *C. aquatica* (**Fig. 2C**). Therefore, when animals are switched from *E. coli* to pathogens, a mixed transcriptional response involving regulation of metabolic and immune response is triggered. However, in the long term, the response is specified and reduced to a small subset of downregulated genes that are common between pathogens and nutritious food (**Fig. S1C**), showing that the transcriptional response prior to dauer formation in the F2 reflects a defensive response to pathogenic conditions intertwined with an ongoing metabolic transformation.

**Fig. 2.**
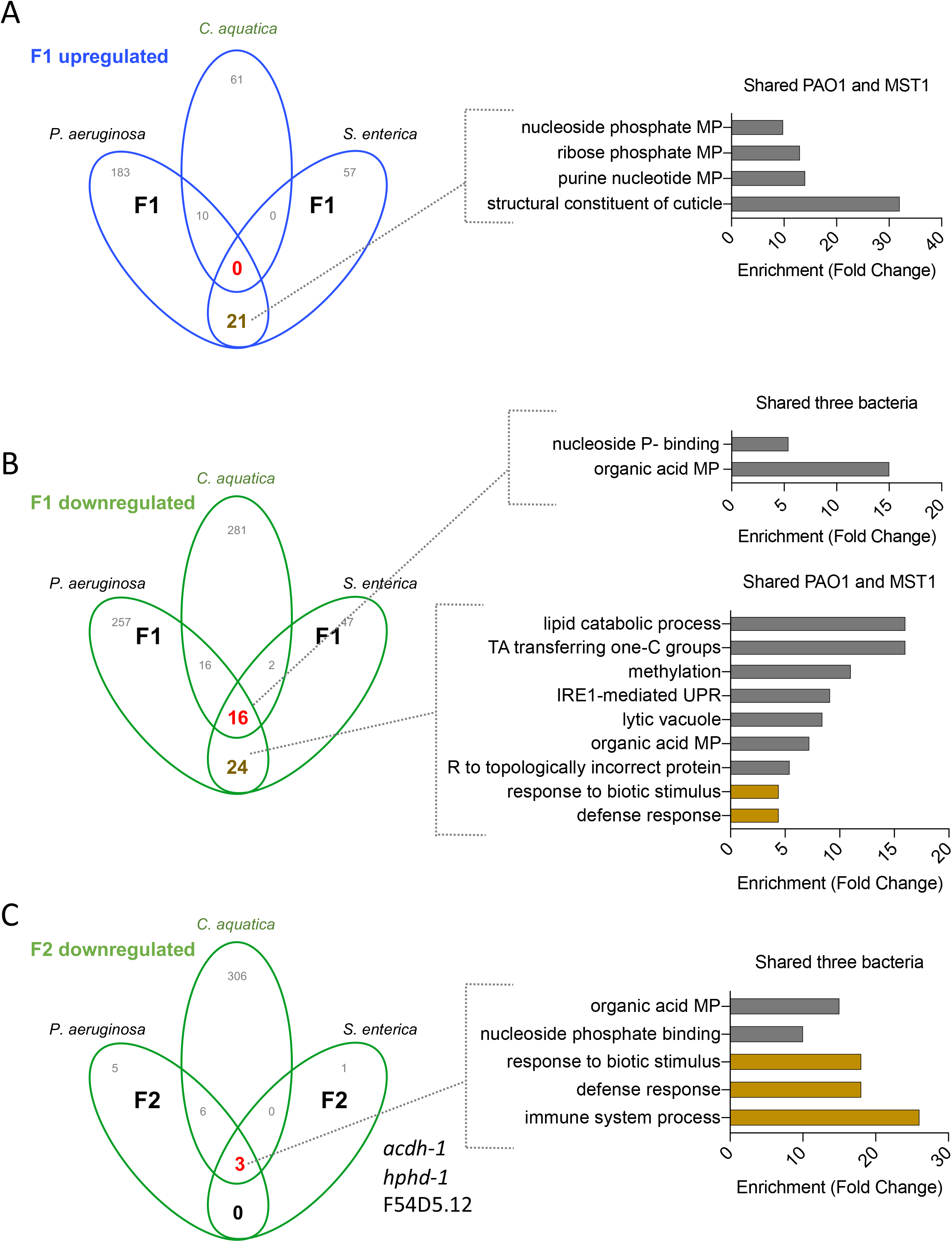
Coincidences in upregulated (A) and downregulated (B-C) genes of animals feeding on pathogens and those reported for *C. aquatica* (14), for one (A-B) and two (C) generations after change from *E. coli* OP50. Each figure includes GO enrichments for genes shared with *C. aquatica* and those shared between pathogens.

In addition, to relate transcriptional changes exclusively induced by pathogen exposure to processes of physiological relevance for PIDF, we examined enriched GO terms in up and downregulated genes, in each generation and each bacterium (**Fig. S1A-B** and **D-E**). Upregulated genes in the F1 on both pathogens shared enrichment in structural components of the cuticle (Fig. **S1A-C**). *S*. *enterica* could only be tested in the F1 because there was only one differentially expressed gene in the F2. In *P. aeruginosa*, both generations displayed GO enrichment, with the F2 specifically enriched in genes involved in defense against biotic stress (**Fig. S1A** and **C**). The latter was significantly and specifically enriched in animals fed on *P. aeruginosa* for two generations (**Fig. S1D-F)**. Taken together, these results further prompt the idea that in the long-term, *C. elegans* can adapt to the pathogenic encounter overcoming the general response to diet change and keep specific biological changes that may aid their survival, such as cuticle, metabolism, defense and dauer reprogramming.

Global analysis of mRNA overexpression shows that changes are dissimilar for animals feeding on *S*. *enterica* serovar Typhimurium MST1 and *P. aeruginosa* PAO1 throughout the two generations. Since worms can enter diapause in both *P. aeruginosa* and *S. enterica* the decision that gives rise to PIDF is not solely reliant on changes in mRNA expression, but also on other levels of regulation. To further understand its underlying molecular causes, we analyzed changes in the sRNAs repertoire in both bacteria and generations.

### sRNA expression in two generations of *C. elegans* exposed to bacterial pathogens

Small RNAs (sRNAs) are broad regulators of gene expression (15, 16, 17) and key candidates to modulate inter and transgenerational environmental adaptation (13, 18, 19). We aimed to unbiasedly identify known and novel sRNAs expressed when animals are fed on each bacterium through two consecutive generations. For that, we defined candidate sRNAs loci based on transcriptional peaks coordinates or TPs (defined in Materials and Methods). These TPs were used for the downstream RNAseq analysis. Then we compared those genomic loci with annotated features to classify them as known (matching an annotated sRNA), novel (located in intergenic regions), or partially novel (unannotated but overlapping or nested within a feature, see **Fig. 3A** and **Dataset 2**). Considering all differentially expressed genes in either pathogen 6.2% were known features, 77.8% were partially novel sequences, and 16% were novel TPs (**Fig. 3B**). Known differentially expressed TPs in pathogen-fed worms include pre-miRNAs and mature miRNAs. Partially novel TPs were nested in intronic or exonic segments of coding genes, or overlapping with 5’ and 3’UTR ends. The TPs nested in non-coding transcripts were found in rRNAs and 21-ur RNAs. We also found pseudogenic transcripts, tRNAs and novel TPs within intergenic regions (**Fig. 3C** and **Table S4** and **5**). Interestingly, from all expressed TPs, *mir-243*-3p along with another 11 TPs nested within the *rrn-3.1* ribosomal gene were upregulated in both generations of worms fed on *P. aeruginosa* PAO1 and *S*. *enterica* MST1 (**Fig. 3D** and **Table S6**). In contrast, the pre*mir-70* and another 9 novels TPs were downregulated similarly in both, bacteria and generations (**Fig. 3E** and **Table S6**, shared all conditions). Despite these, most TPs were specifically DE in response to MST1 or PAO1, or to both pathogens but in only one generation. For example, *mir-51* was exclusively upregulated in the F1 of animals feeding on PAO1 (**Table S6**, generation-specific), but *21ur-6043* was upregulated in both generations on PAO1 (**Table S6,** bacterium-specific). Interestingly, 4 miRNAs (*mir-1, mir-48, mir-256, mir-257*) were downregulated only in the second generation on *P. aeruginosa* PAO1 (**Table S4**). One of them, *mir-48*, a *let-7* Fam member is known to be repressed on the more virulent *P. aeruginosa* strain PA14 (20). These results show that sRNA expression in the context of long-term pathogen exposure is mostly specific to one generation or one pathogen. However, *mir-243-3p* is overexpressed across conditions suggesting it may function as a common effector in PIDF. As the response to PAO1 in the F2 was the largest in terms of numbers of TPs both up and downregulated (**Fig. 3F**) the following experiments were performed using *P. aeruginosa* PAO1.

**Fig. 3.**
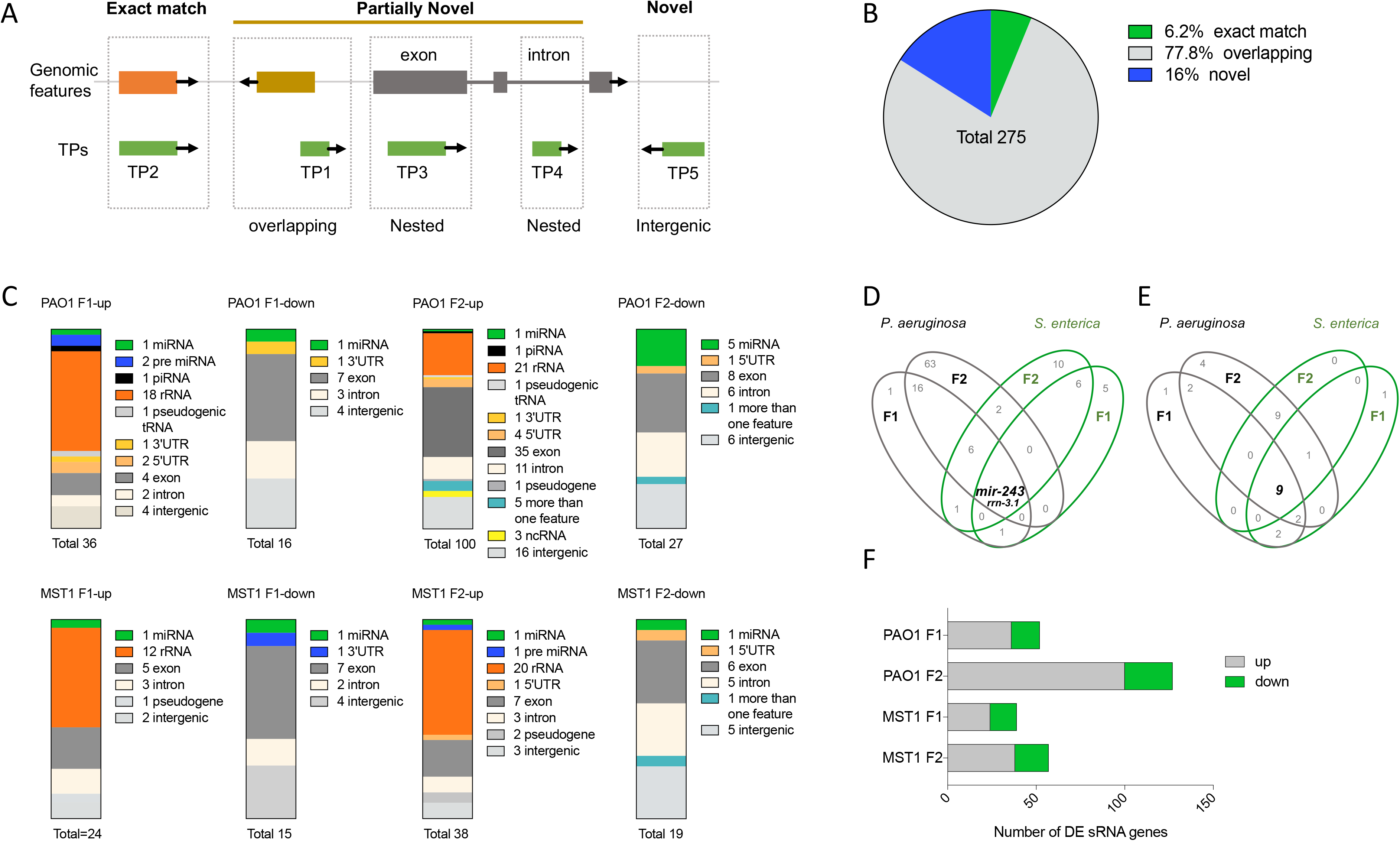
Global analysis of differential small RNA gene expression of an intergenerational infection paradigm **A**. Representation of transcriptional unit designation. **B**. Genomic context of sRNA genes differentially expressed in pathogenic conditions. **C.** Type and abundance of sRNA genes differentially expressed on *P. aeruginosa* PAO1 and *S*. *enterica* MST1 in two generations **D-E**. Venn diagram representation of shared and unique genes over-expressed (D) and repressed (E) in each generation and on each pathogen. **F**. Number of sRNA genes differentially expressed on *P. aeruginosa* PAO1 and *S. enterica* MST1 in two generations.

### *mir-243* is necessary for diapause formation under pathogenesis

To test the requirement of *mir-*243 on PIDF we first quantified by RT-PCR the relative amounts of mature *mir-243-3p* in each generation of animals feeding on *P. aeruginosa* PAO1 compared to those feeding on *E. coli* OP50. RNA was extracted from L2 worms fed on *P. aeruginosa* and *E. coli* in the F1 and F2, as was done for the transcriptomic analysis. We found that *mir-243-3p* appeared upregulated by 4-fold in the second generation of worms feeding on *P. aeruginosa* PAO1 and 1-fold change in the F1, and as expected, *mir-243* (*n4759*) mutants were unable to express mature *mir-243* (**Fig. 4A**). To further explore the role of *mir-243* in PIDF we tested whether mutant animals for *mir-243* (*n4759)* were able to form dauers on *P. aeruginosa* PAO1 in the second generation. We also tested as reference animals with a deletion in *mir-235 (n4504)*, a microRNA involved in nutritional-related L1 diapause (21), which was not DE under pathogenesis. *mir-243* and *mir-235* mutants were fed on *P. aeruginosa* PAO1 for two generations. The population growth and appearance of dauers was quantified in the F2 and compared to wild type animals. Growth on *P. aeruginosa* PAO1 was not affected by either mutation (**Fig. 4B**), but just *mir-243* mutants were found to form significantly less dauers than wild type or *mir-235* animals when fed on *P. aeruginosa* PAO1 (**Fig. 4C**). Therefore, these mutations do not increase susceptibility to pathogens, as revealed by growth, but since none of them were deficient in dauer formation under starvation (**Fig. 4D**), this result importantly suggests that *mir-243* has a specific role in dauer formation under pathogenesis.

**Fig. 4.**
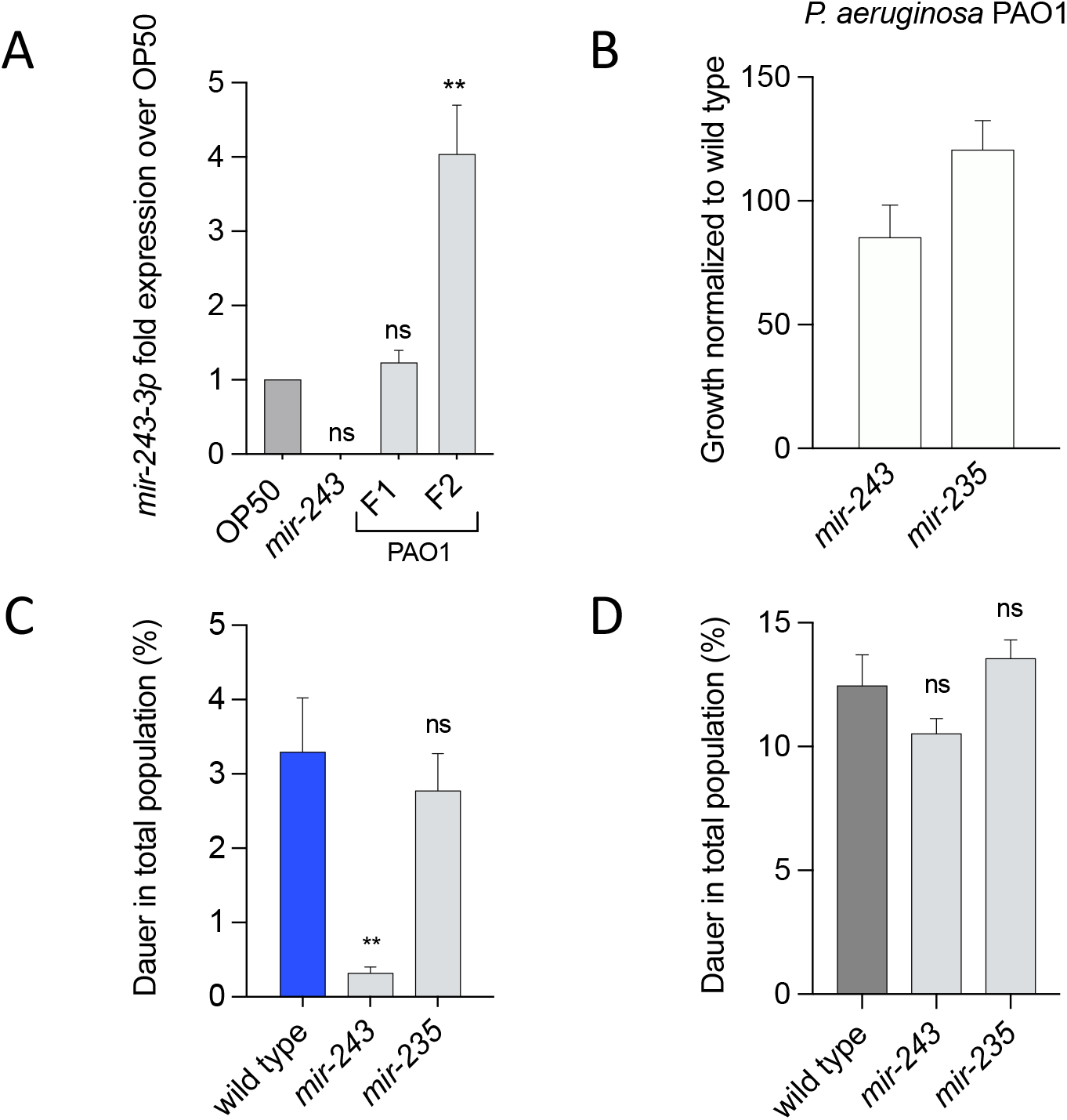
Expression of mature *mir-243* and role on pathogen induced diapause. **A**. Quantification of *mir-243-3p* expression in animals feeding on *P. aeruginosa* PAO1 for two generations. **B-D**. Growth (B), dauer formation on pathogens (C) and dauer formation by starvation (D) of *mir-243* mutant animals.

We have shown that *mir-243* is required for PIDF and thus we explored its potential molecular targets. Interestingly, *mir-243* is known to activate the exo-RNAi pathway by binding RDE-1, triggering the production of secondary siRNAs. the Y47H10A.5 mRNA has been revealed repressed by this mechanism (20). Even though *mir-243 i*s upregulated in all PIDF conditions, Y47H10A.5 is not differentially expressed in our data. Therefore, we tested the hypothesis that some downregulated mRNA under pathogenesis could be *mir-243* targets. Using the IntaRNA tool version 2.0 (22), we computed the expected RNA-RNA interactions among *mir-243* and our downregulated mRNAs. We found that *mir-243* has the potential for binding 26 of our 136-downregulated genes with high complementarity (seed >=12) and strong negative free energy (NFE). Even though Y47H10A.5, a validated *mir-243-3p* target, is not differentially expressed in our data, we computed the interaction with the same parameters and found that Y47H10A.5 has lower NFE and shorter seed that our candidate targets (**Table S7**).

### DAF-16 and other transcriptional activators regulate the expression of *mir-243* under pathogenesis

To study whether the increase of mature *mir-243* in animals feeding on pathogens in the F2 could be a result of transcriptional activation we quantified the fluorescence of a strain expressing *gfp* under the *mir-243* promoter (VT1474). We measured *gfp* expression in L2 worms grown on pathogens for two generations and compared it with those grown on *E. coli* OP50. Animals feeding on pathogens overexpress *Pmir-243::gfp* compared to *E. coli* OP50 controls in both generations (**Fig. 5A**) suggesting that the exposure to pathogens activates the transcription of *mir-243* in concordance with our previous expression results (**Fig. 4A**). However, the mechanism behind *mir-243* transcriptional activation is unknown. A number of transcription factors (TF) promote the expression of their targets under stress and infection (23, 24, 25). It has been previously reported that *mir-243* and members of the *let-7*fam are the miRNAs with the highest number of interactions to TF in the *C. elegans* genome (26). We specifically tested the role of three transcriptional regulators, DAF-16, PQM-1 and CRH-2 in *mir-243-3p* expression. The reasons for choosing them are explained below. In our paradigm, DAF-16/FOXO transcriptional activator localizes into the nucleus of animals exposed to *P. aeruginosa* PAO1 prior to diapause formation (13). PQM-1 resides in the nucleus regulating the expression of DAF-16-associated elements (DAE), avoiding dauer formation (27). Finally, we included CRH-2 because is a direct target negatively regulated by the *let-7* family of microRNAs (28), which have been involved in the response to pathogenesis by the *P. aeruginosa* PA14 strain (20). In our experiments, *mir-48*, a *let-7*fam miRNA, was downregulated in PAO1 in the F2 (**Table S4**). Under this logic, *crh-2* could be indirectly upregulated through *mir-48* downregulation. To give further ground to this selection, we tested whether *pqm-1* and *crh-2* promoters expression were higher in pathogens compared to non-pathogenic conditions. Using strains expressing *gfp* under promoters for *pqm-1* or *crh-2* we observed that the expression of both TFs is upregulated in animals fed with *P. aeruginosa* PAO1 for two generations compared to those fed on *E. coli* OP50 (**Fig. 5B** and **C**), as we have previously reported for DAF-16 (13). To study whether these transcriptional regulators are necessary for the expression of *mir-243* under pathogenesis we extracted RNA from *daf-16 (m27)*, *pqm-1 (ok485)* and *crh-2(gk3293)* mutants in the F2 of worms fed with *P. aeruginosa* PAO1 and quantified the expression of *mir-243-3p* over wild type animals. All three mutants fed on *P. aeruginosa* for two generations completely lacked *mir-243-3p* expression (**Fig. 5D**). These results show that DAF-16, PQM-1 and CRH-2 transcription factors are needed for the expression of *mir-243-3p* in the second generation of animals exposed to pathogens.

**Fig. 5.**
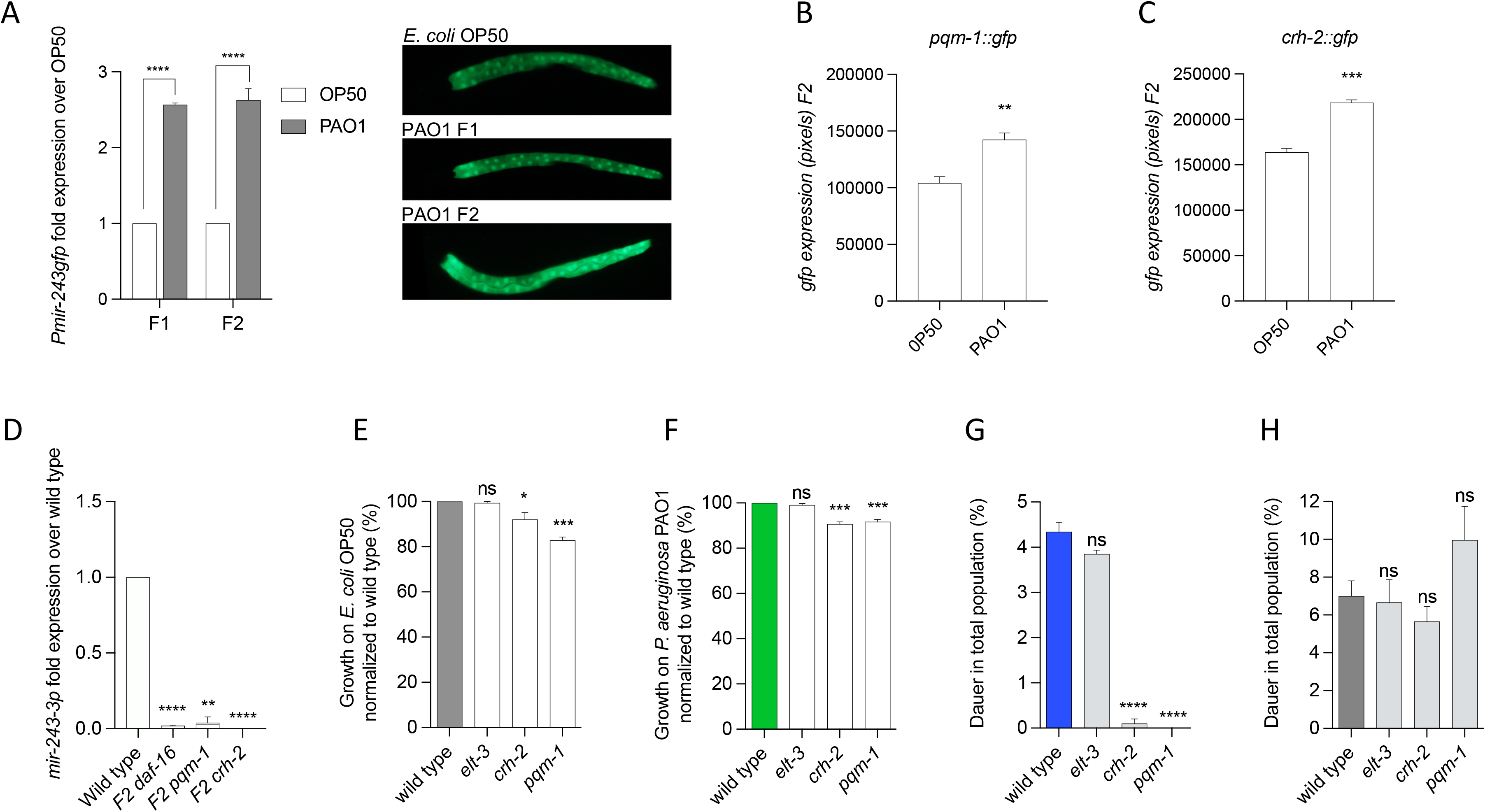
Transcriptional factors required for *mir-243* expression and pathogen-induced diapause formation. **A**. Quantification of *mir-243* promoter expression by GFP in animals feeding on *P. aeruginosa* PAO1 compared to *E. coli* OP50 and representative photos. **B-C**. Quantification of expression of PQM-1 (B) and CRH-2 (C) by GFP expression on animals fed on pathogens. **D**. Quantification by RT-PCR of *mir-243-3p* on wild type and *daf-16*, *pqm-1* and *crh-2* mutant animals. **E-G**. Growth on *E. coli* OP50 (E), *P. aeruginosa* PAO1 (F) and dauer formation (G) in the second generation of animals. **H**. Dauer formation on starvation of wild type and mutants of transcription factors.

Because *mir-243* loss affects the ability of animals to enter diapause under pathogenesis we further explore whether *crh-2* and *pqm-1* mutants also failed to form dauers under infection but formed normal amounts of dauers under starvation. Additionally, we tested whether these mutants were able to grow on *E. coli* and *P. aeruginosa* to wild type extents. As a control, we used a mutant of *elt-3*, a transcriptional activator with a broad expression in the animal. *daf-16* mutation causes animals to be unable to form dauers (29) and could not be tested. All TF mutants grew well on *E. coli* OP50 and to a similar extent as wild type animals on pathogenic bacteria (**Fig. 5E** and **F**). Interestingly, *crh-2* and *pqm-1* mutants were unable to enter diapause under pathogenesis after two generations (**Fig. 5G**) but formed normal amounts of dauers under starvation (**Fig. 5H**), while *elt-3* animals formed normal amounts under both, pathogenesis and starvation (**Fig. 5G** and **H**). Taken all together these results suggest that the role of *crh-2* and *pqm-1* TFs is specific to PIDF, and that an expression signaling cascade including CRH-2, PQM-1 and DAF-16 upstream of *mir-243* expression is triggered by the long-term interaction of worms with the mild pathogen *P. aeruginosa* PAO1.

## Discussion

Survival strategies to cope with environmental challenges rely on the genetic plasticity of organisms. In this work, we dissected the transcriptomic differences and similarities between worms feeding on three different bacteria. Two of them, *P. aeruginosa* PAO1 and *S*. *enterica* serovar Typhimurium MST1 elicit dauer entry as a defense strategy in the second generation (13). Differential gene expression analysis allowed us to identify *mir-243-3p* (the mature form of *mir-243*) as the only common upregulated sRNA in animals fed on these pathogens for two generations. Moreover, mutant animals for *mir-243* do not perform PIDF despite not being dauer defective under starvation. Finally, we tested the role of transcription factors DAF-16, PQM-1, and CRH-2 on *mir-243-3p* expression under pathogenesis. All three showed to be required for *mir-243* expression. Furthermore, in contrast to dauer defective DAF-16, the CRH-2 and PQM-1 transcription factors are specifically required for PIDF but not for dauer formation under starvation.

### Most transcriptional changes are unique to the encounter with each pathogen

Gene expression is highly variable and dependent on environmental and physiological factors. *C. elegans* transcriptional profile is modified when animals are exposed to a new bacterial diet. These can be nutritious food, as *Comamonas aquatica* (14), non-pathogenic such as *Bacillus subtilis* (30), or pathogenic such as *Shigella flexneri* (31), *P. aeruginosa* PA14 and *Staphylococcus aureus* (32, 33). Temperature and diet changes trigger gene expression changes associated with defensive responses and metabolism (30). Environmental conditions can change the phenotype of subsequent generations (34, 35, 36) suggesting that transcriptomic modifications can be inherited to the progeny. Accordingly, we speculate that the greater abundance in polyA+ transcripts on the F1 may reflect the response to a novel food source (being their first time not feeding on *E. coli* OP50, **Fig. 2**) and generalized stress and immune responses to pathogenesis. Therefore, the second generation of worms forced to fed on pathogens may have adapted to the pathogenic bacteria through inherited signals, narrowing the transcriptional changes to a more reduced transcriptional response [**Fig. S1**, (37)].

In this work, we discovered transcriptional changes that could explain the defensive decision to enter the dauer program as a response to pathogenesis (13). We notice that two different bacterial pathogens trigger the same phenotypic response in *C. elegans* but display few underlying transcriptomic similarities. We think that this could be an indicator of high functional specificity, result from a finely tuned long-term communication between bacteria and host. Clustering of genes by function (38) revealed large differences between animals fed on the two bacteria. In the wild, *C. elegans* is mostly found in the dauer stage, a strategy used to maximize the animal’s survival by ensuring their dispersal to new food sources (39). Dauer entry, therefore, may be a convergent phenotypic outcome driven by a plethora of different stimulus and transcriptional regulatory pathways that allow the animal, under adverse circumstances, to ensure their own specie’s survival.

### Small RNAs and pathogenesis

Several works have explored the role of sRNA in infection (40, 41). Among them, microRNAs have been reported to be involved in the innate immune response of *C. elegans* against infection with bacterial pathogens, and even with eukaryotes such as the opportunist yeast *Candida albicans* (42). For example, *let-7* regulates the innate immune response by targeting intestinal SDZ-24 when fed on the highly pathogenic *P. aeruginosa* PA14 (43). Moreover, other candidate targets of *let-7* and *let-7*Fam of microRNAs may include components of the PMK-1/p38 innate immune pathway (20). Likewise, *mir-67* mutants exhibited reduced pathogen avoidance behavior, apparently due to a dysregulation in *sax-7* targeting (44, 45). Others like *mir-70* and *mir-251/mir-252* mutants possess enhanced survival to *P. aeruginosa* PA14 infections, indicating that these miRNAs negatively regulate the immune response (46). Supporting this idea, we found that *mir-70* was systematically downregulated in animals feeding on both *P. aeruginosa* and *S. enterica*.

We describe here, that a single sRNA, *mir-243*, was the only upregulated transcriptional coincidence among animals fed on pathogens for two generations, and resulted to be required for pathogen-induced dauer formation. As mentioned, *mir-243* has an unusual association with RDE-1, an Argonaut protein known to be a siRNA acceptor (47), suggesting that *mir-243* may induce mRNA destabilization depending on the RNAi machinery. In accordance with that, we have previously shown that RDE-1, along with other RNAi effectors, are needed for the induction of dauer formation under pathogenesis (13). Predicted targets of *mir-243* within downregulated genes were almost perfectly complementary, suggesting that the mechanism by which *mir-243* induces silencing in the context of PIDF is could be also related to siRNA pathways as previously reported for other targets (47). Our approach allows us to narrow the spectra of possible *mir-243* targets in our experimental paradigm. *mir-243* can potentially target many genes for silencing. Côrrea et al. (47) reported 1835 upregulated genes in the *mir-243* mutant (microarray analysis comparing adult wild-type and mutant worms, fold change >= 2, p < 0.05). We found 24 common genes between published upregulated genes in *mir-243* mutant (47) and downregulated genes in our datasets. Three of them (C14B1.3, *acs-2* and *mrp-2*) have high negative free energy (NFE) of interaction and high sequence complementarity (> or = 12 bp), becoming good candidates for future validation studies. The biological validation of predicted *mir-243* targets, as well as the role of downregulated microRNAs, such as *mir-70*, in PIDF will remain unresolved. Notwithstanding, our findings support the hypothesis that the response of *C. elegans* to different pathogens is accompanied by dynamic changes in the activity of miRNAs. In this work, we found that *mir-243* is involved in the response of *C. elegans* to infection with both pathogens, *P. aeruginosa* PAO1 and *S. enterica* MST1. Moreover, as the same miRNA is triggering similar effects on different pathogens, this may imply that changes in a single microRNA can induce similar phenotypic outputs, like dauer formation, but distinct molecular cascades to achieve it.

### Transcription factors involved in diapause formation as a defensive strategy

A number of transcription factors activate the transcription of their targets under stress and infection. DAF-16 has been reported to regulate the expression of genes involved in defense when exposed to pathogenic bacteria such as *P. aeruginosa* PA14 (48), *S. enterica* strain 1344, *Yersinia pestis* strain KIM5, and *Staphylococcus aureus* MSSA476 (49). In the pathogen-induced dauer formation paradigm, DAF-16 localizes in the nucleus of animals exposed to *P. aeruginosa* PAO1 prior to diapause formation (13). In this work, we show that *mir-243* expression requires intact DAF-16. Furthermore, both, PQM-1, that affects the expression of DAF-16 in the nucleus (27), and CRH-2, that has been proposed to be regulated by *let-7* miRNA (28), are needed for *mir-243* expression under pathogenesis and their loss impairs PIDF. Although we did not carry out an exhaustive analysis of TFs related to the formation of dauer by pathogens, an extensive list of TFs that interact with *mir-243* and that could regulate it directly or indirectly, is available in the work of Martinez et al. (26). In this work we did not find differentially expressed genes whose mutations have been described before as conducing to abnormal dauer formation (*Daf*). This suggests at least two things, the expression of *Daf* genes does not necessarily change at the level of mRNAs, or that the dauer program under pathogenesis is molecularly different from starvation-induced diapause (an abiotic stressor).

Dauer formation upon pathogenesis is likely a multistep process that involves the sensing, initiation and establishment of the pathogenic state. The transcriptional difference in polyA+ genes of animals feeding on either pathogen during the F1 is much larger than in the F2 compared to their usual *E. coli* OP50 food. We speculate that the F1 response is pleiotropic and involves i) the first encounter with new bacteria and diet, and ii) the response to a pathogen. Importantly, the worm expression profile is mostly specific for each bacterium, coherent with the dissimilar nature of *P. aeruginosa* and *S. enterica* (50). This polyA+ response is dramatically reduced in the F2, where differentially expressed genes are specifically involved in immune and defense response. Therefore, it is likely possible that in the second generation, specific transcriptional changes in polyA+ coding and non-coding genes are accumulated sufficiently to exceed the threshold thus modulating developmental decisions to ensure survival. In this context, miRNAs and other sRNAs play key regulatory roles required for phenotypic and inter/transgenerational responses in *C. elegans.*

Dauer entry is a hard decision because it is metabolically and reproductively expensive for the animal. Therefore, we presumed that dauer-triggering signals should exceed a threshold that supports this choice. Time-dependent signaling molecules may be insufficient to reach a temporal threshold in the first generation of animals exposed to pathogens. For example, we know that dauer entry requires persistent intestinal colonization, which takes more than 48 hours feeding on *Pseudomonas aeruginosa*. Dauer entry is a complex process in which immune, metabolic, and stress signals are integrated at different levels of regulation.

## Materials and Methods

### *C. elegans* and bacterial growth

Wild type, mutant and transgenic *C. elegans strains* were grown at 20°C as previously described (51). All nematode strains were grown on *Escherichia coli* OP50-1 (resistant to streptomycin) before pathogen exposure. *S. enterica* serovar Typhimurium MST1 (ATCC 140828) and *P. aeruginosa* PAO1 (ATCC 15692) were used for infection protocols. All bacteria were grown overnight on Luria-Bertani (LB) plates at 37°C from glycerol stocks. The next morning a large amount of the bacterial lawn is inoculated in LB broth and grown for 6 hours at 250 rpm and at 37°C. 3 mL of the resulting bacterial culture is seeded onto 90 mm NGM plates and allowed to dry for 36 hours before worms are placed on them.

### *C. elegans* strains

We used the following strains of the *Caenorhabditis* Genetics Center (CGC): Wild type (N2), MT15454 [*mir-243*(*n4759*)], MT16060 [*mir-253*(*nDf64*)], DR27 [*daf-16*(*m27*)], VC3149 [*crh-2* (*gk3293*)], RB711 [*pqm-1* (*ok485*)], VC143 [*elt-3* (*gk121*)], and transgenic strains VT1474 [*Pmir-243::gfp* (*unc-119* (*ed3*) III; *maIs177*)], OP201 [*unc-119* (*tm4063)* III; wgls201 (*pqm-1::TY1::EGFP*)], BC13136 [*crh-2* C27D5.4a*::gfp (dpy-5* (*e907*); sEx13136)], BC14266 [*dpy-5* (*e907*); sEx14266 (rCesF35E8.8::*GFP +* pCeH361)]. Pertinence of strains with mutations in TF genes: There are many available strains with mutations in *daf-16*. We chose DR27 because it was the strains with the strongest Daf-d phenotype under starvation. *pqm-1(ok485)* and *crh-2(gk3293)* are the only strains available in the CGC for those genes.

### Hypochlorite treatment

To synchronize *C. elegans* and/or to obtain pure embryos, we prepared a 5% hypochlorite solution containing 20 mL of 1 M NaOH, 30 mL NaClO and 50 mL H_2_O in 100 mL final volume. Plates with mostly gravid adults were washed with 1 mL of M9 (KH_2_PO_4_ 3 g, Na_2_HPO_4_ 6 g, NaCl 5 g, 1 M MgSO_4_, H2O up to 1 L) and transferred to an Eppendorf tube. The volume collected was centrifuged at 394 × g. 1 mL of the hypochlorite solution was added to the pellet. After 5 minutes of vigorous vortex, the tube was centrifuged at 394 × g for 2 minutes. The pellet was washed with 1 mL of M9 solution and centrifuged at 394 × g and 2 minutes to discard the supernatant. The resulting pellet contains an embryo concentrate.

### Quantification of population and dauer larvae

*Dauer formation on pathogens.* Entire worm populations on each plate were collected in 1 mL of M9. This initial stock was diluted 1:10 in M9. 10 μL of this 1:10 dilution was used to count the total population of worms under a Nikon SMZ745 stereomicroscope. To quantify the number of dauers in each population, the initial stock was diluted 1:10 in a 1% SDS solution and maintained in constant agitation for 20 minutes (52). To count the number of total animals and dauers, 10 μL of this last dilution was placed in a glass slide under the stereomicroscope. Each condition was scored 3 times (triplicated of one technical replica) and dauers were plotted as a percentage of the total populations of animals.

### *C. elegans growth* in pathogenic bacteria

Five L4 (P0) wild type worms or mutants (grown in *E. coli* OP50) were picked and transferred to a 90 mm diameter plate seeded with 3 mL of *P. aeruginosa* PAO1, *S. enterica* serovar Typhimurium MST1 or *E. coli* OP50 control bacteria. In all cases, the bacterial lawn covered the plate. After 24 hours the F1 embryos were obtained by hypochlorite treatment. All obtained embryos were placed on a new plate with *P. aeruginosa* PAO1, *S. enterica* serovar Typhimurium MST1 or *E. coli* OP50. Animals were allowed to grow for 24 hours until they reached the L2 stage. The total number of worms in the population and the percentage of dauer were quantified.

To obtain the F2, five L4 larvae were transferred from *E. coli* OP50 to a 90 mm diameter plate with 3 mL of *P. aeruginosa* PAO1 or *S. enterica* serovar Typhimurium MST1. After eight days the total number of worms and dauer larvae were quantified. The number of bacteria seeded allowed animals to be well fed for the length of the experiment. In the case worms starved, we discarded that experiment. Each assay was performed in three independent experiments (technical replicates) generating a biological replica. A total of three biological replicates were considered for each analysis.

### Dual RNA Seq

#### Sample preparation

Wild type *C. elegans* were cultured on 60 mm diameter Petri dishes with NGM media seeded with 500 μL of *E. coli* OP50 and maintained at 20°C. After three days, mixed stage animals were treated with bleaching solution and embryos were deposited in new dishes. 48 hours later, most individuals were in the L4 stage. Five L4 worms were transferred to 90 mm plates previously seeded with 3 mL of *E. coli* OP50, *S. enterica* serovar Typhimurium MST1 or *P. aeruginosa* PAO1. In all cases, the bacterial lawn covered the plate. Worms were allowed to grow at 20°C for 24 hours. After that time, all animals on the plate were subject to hypochlorite treatment. F1 embryos were collected in 1 mL of M9 and centrifuged at 394 x g. The embryos obtained were placed on a 90 mm new plate with 3 mL of bacteria. After 24 hours, animals were collected with M9 for total RNA extraction. Worms on other 3 plates were allowed to grow for another 48 hours until the F1 was gravid. F2 progenies were collected in the same way as the F1 and placed on a separate plate with the same species of bacteria. Animals were collected for RNA extraction 24 hours later. Each condition was performed in triplicates obtaining a total of 18 samples (F1 and F2 in *E. coli* OP50, *P. aeruginosa* PAO1 or *S. enterica* serovar Typhimurium MST1).

### RNA extraction

#### RNA extraction of colonizing bacteria and C. elegans

Worms were washed off the plates with 1mL of M9, centrifuged at 394 × g for 2 min and resuspended at least 5 times with M9. RNA purification was performed using TRIzol (Life Technologies) following manufacturer’s instructions. For RNA extraction, samples were mechanically lysed by vortexing with 4 mm steel beads for no more than five minutes. Each condition was performed in triplicates. Several biological replicas were mixed to reach the required concentration of 1 μg/μl of total RNA.

#### mRNA library preparation and sequencing

Total RNA was isolated from synchronized F1 and F2 *C. elegans* populations feeding on non-pathogenic *E. coli* OP50, and pathogens *P. aeruginosa* PAO1 and *S*. *enterica* serovar Typhimurium MST1 as explained above. After DNase I digestion with DNAse I (Invitrogen™), RNA concentration was measured using Quant-iT™ RiboGreen^®^ RNA Assay Kit (Life Technologies). The integrity of RNA was determined on the Agilent 2100 Bioanalyzer (Agilent Technologies). mRNA libraries were prepared with the IlluminaTruSeq™ RNA sample preparation kit (Illumina) according to the manufacturer’s protocol. The quality and size distribution of the libraries were evaluated with the Agilent 2100 Bioanalyzer using a DNA 1000 chip (Agilent Technologies) and quantified using the KAPA Library Quantification Kit for Illumina Platforms (Kapa Biosystems) on the Step One Plus Real-Time PCR System (Applied Biosystems).

The *C. elegans* mRNAs libraries were sequenced using the HiSeq Illumina platform (BGI) with paired-end sequencing (2x 100 bp, BGI). *C. elegans* and bacterial small RNA were sequenced using the Mi-Seq Illumina platform at the Center for Genomics and Bioinformatics (CGB), Universidad Mayor.

#### sRNA library construction and sequencing

Samples were prepared and sequenced at CGB. Libraries were constructed with the TruSeq sRNA Sample Preparation Kit (Illumina), according to the manufacturer’s instructions. For quality control, cDNA libraries were run on a high sensitivity DNA chip using the Agilent 2100 Bioanalyzer (Agilent Technologies), according to the manufacturer’s instructions. An agarose gel library size selection was performed to recover RNAs shorter than 200 nucleotides. The library was quantified using a high sensitivity DNA chip. The sRNA libraries were sequenced in an Illumina MiSeq sequencer using the MiSeq Reagent v2 50-cycle kit with single-end sequencing (1 × 36 bp). For each biological condition, three libraries were constructed from biological replicates.

### Bioinformatic analysis

#### mRNA transcriptomics of C. elegans

##### Data pre-processing and quality control

Trimming was made with Trimmomatic v. 0.36 (53). Reads with a quality score (Phred score) less than 35 and read length less than 36 were removed.

##### Mapping and read count

Reads were aligned using Tophat (54) with default parameters. Reads were quantified using HTSeq count (55).

*Differential expression* was determined using EdgeR (56) and DeSeq (57). Differentially expressed genes were defined as those with adjusted p-value (padj) <0.05 by either method.

*Enrichment analysis: Gene Ontology (GO) analysis* was performed using the enrichment tool in wormbase (58).

#### Small RNAs transcriptomics of C. elegans

##### Data pre-processing and quality control

Quality visualization was made with FastQC (http://www.bioinformatics.babraham.ac.uk/projects/fastqc). Illumina small 3’ adaptors were removed with Cutadapt version 1.13 (59). Reads with an average quality over four bases lower than 30, as well as reads shorter than 17 bp were discarded with trimmomatics (53).

##### Mapping

For each sample, the reads were aligned against the *C. elegans* genome PRJNA13758 WS267 using bowtie2 version 2.2.6 (60). We chose a 17 bp seed length, which was the length of the shorter read. The seed interval was calculated as f(x) = 0 + 2.5 * sqrt(x), where x is the read length (roughly two seeds per read as reads are 36 or smaller). For greater sensitivity, we set the number of mismatches allowed in a seed to 1. By default, 15% of ambiguous characters per read is allowed. A bam file was produced for each sample.

##### Defining features and counting reads

We developed a strategy to detect both known and unannotated transcripts from bacteria and *C. elegans.* For that, we chose to make a reference annotation based on observed expression peaks. Some reads overlap annotated genes, others show more than one peak in annotated regions, and many peaks are located in unannotated regions. We selected those genomic areas covered by mapped reads to define customized features, named transcriptional peaks (TPs) defined as a genomic area that shows a peak of expression (more than 10 per base) and that may correspond to direct transcription or fragments of a longer transcript. This was done by merging bam files from each sample using SAMTools version 0.1.19 (61). Afterward we only kept features between 17 and 150 nucleotides with an average coverage of 10 or more reads by nucleotide. The features obtained for both strands were gathered and sorted to create a custom GFF file for further analysis. Next, we counted the reads against those custom GFF. For comparison with databases, we intersected TPs with reported annotations and classified them according to their genomic context.

##### Comparison with annotated genes

To see how TPs matched with annotated genes, we compared them to the Ensembl database, by intersecting the GTF file of *C. elegans (PRJNA13758.WS267)* with our custom GFF file, using BEDTools (62). Based on this result, we classified TPs as novel (in intergenic unannotated regions), nested or overlapping annotated features, and sense or antisense to a known feature.

##### Differential expression analysis between conditions

For each sample, read count was performed with featureCounts (63) from the Bioconductor Rsubread package with default parameters. Then, the count matrix was used to perform differential expression analysis in R (version 3.3.2) between worms fed with three different bacteria of each generation using DeSeq2 (64) version 1.4.5. The samples of animals feeding on pathogenic strains *P. aeruginosa* PAO1 and *S. enterica* serovar Typhimurium were compared with *E. coli* OP50. For each condition, the first and second generations were compared.

We conducted the previous analysis using a custom-made bash and R scripts, available at https://github.com/mlegue/Gabaldon_2020.

##### Prediction of *mir-243-3p* targets

We used the IntaRNA tool version 2.0 (22); http://rna.informatik.uni-freiburg.de/IntaRNA/Input.jsp. We adjusted parameters to force the longest possible seed with perfect complementarity. We started using the maximal admitted seed length (SeedBP=20) and run the analysis with all seed length above default (SeedBP=8) with no restrictions on seed energy. We also set the temperature to 20°C at which experiments were performed. We performed the interactions between *mir-243*-*3p* and the downregulated mRNA genes (polyA-genes) in our dataset. We also evaluate with the same parameters the interaction with Y47H10A.5, a validated *mir-243-3p* target.

### Quantification of differential expression *by RT-PCR*

#### Total RNA Extraction

5 L4 (P0) worms were placed in 90 mm NGM plates seeded with *P. aeruginosa* PAO1. After 24 hours, the total population was collected with 1 mL of M9 and centrifuged at 394 × g for 2 minutes. The pellet was treated with hypochlorite solution (see above). All resulting F1 embryos were placed in new plate with *P. aeruginosa* PAO1 for 24 hours and collected as L2. Collection is done with M9 in an Eppendorf tube. Contents were centrifuged at 394 × g for 2 min. L2 pellet was washed 5 times with M9 to eliminate the most bacteria from the sample. The pellet of synchronized worms (L2) was used for RNA extraction.

F2 animals were obtained from the same L4 P0 fed with *P. aeruginosa* and used to generate the F1 as explained above. F1 embryos were placed in a new 90 mm plate with 3 mL of *P. aeruginosa* PAO1. After 72 hours, the total population was collected with 1 mL of M9 and centrifuged at 394 × g for 2 minutes. The pellet was treated with hypochlorite solution. F2 embryos were placed in a new plate with *P. aeruginosa* for 24 hours. The L2 worms were washed off the plate with M9 and centrifuged at 394 × g for 2 min. 5 washes with M9 were necessary to eliminate most bacteria from the sample. The pellet of synchronized worms (L2) was treated with 4 mm steel beads and 1 mL of RNA-Solv^®^ Reagent (Omega Bio-Tek). The mix was vortexed for 5 minutes. RNA extraction was performed according to manufacturer specifications. Total RNA concentration was quantified with Tecan’s NanoQuant Plate™. Each condition was performed in biological triplicates.

#### cDNA Synthesis

Using the extracted total RNAs, cDNA synthesis was performed with the MIR-X miRNA First Strand Synthesis from TAKARA following manufacturer’s specifications.

#### Real Time PCR (qPCR)

The cDNA samples (concentration 200 ng/μL) were used as a template to perform qPCR with the primers ©QIAGEN Ce_miR-243_1 miScript Primer Assay (MS00019481). The qPCR was performed with the MIR-X miRNA qRT-PCR TB Green Kit from TAKARA. To calculate the relative fold of expression of *mir-243-3p* between generations and genotypes we used delta-delta Ct calculations according to the Reference Unit Mass method (65). This method was based on the comparative use of the test sample against a calibrator (U6). Values less than 1 indicated the negative expression relation with respect to the control and a Ratio greater than 1 indicated the times above the sample with respect to the control.

### Quantification of *gfp* expression

Expression levels of *pqm-1* and *crh-2* genes and the promoter of *mir-243 were* quantified by using wild type animals expressing *gfp*. *gfp* was quantified in F1 and F2 L2 fed with *P. aeruginosa* PAO1 and *E. coli* OP50. Worms were taken individually with a mouth pipette and placed in a bed of agarose with levamisole (20 mM) to immobilize them. To quantify *gfp* expression, we photographed entire animals on a Nikon Eclipse Ni microscope in 40X objective at 1/320s (*mir-243::gfp*) and 1/10s (*pqm-1* and *crh-2*) exposure time, ISO 200/3200 speed (white light and Laser respectively) and focal length of 50 mm. For all markers, photos of entire animals were taken. GFP quantification was done considering the signal from the entire animal. Image analysis was performed using ImageJ. Prior to the analysis images are converted to 8-bit format and a threshold between 1 and 1.2.

### Statistical analysis

Statistical analyzes were carried out using one- or two-way ANOVA with post-hoc tests. For the differential expression analysis, statistical significance was considered lower than p-value <0.05. All experiments were repeated at least three times using technical replicas in each.

---

**Fig. S1.** Enrichment by GO term of upregulated (A, B) and downregulated (D, E) in animals feeding on *P. aeruginosa* PAO1 and *S. enterica* serovar Typhimurium MST1 in two generations. C, F Summary of shared GO terms in F1 and F2 in up (C) and downregulated (F) genes. MP, metabolic process; TA, transferase activity; HA, hydrolase activity; N, Nitrogen; C, Carbon; R, response; P, phosphate; UPR, Unfolded Protein Response.

**Table S1-2**. mRNA coding and non-coding genes differentially expressed in pathogenic conditions. Upregulated (1) and downregulated (2) genes expressed in F1 and F2 of animals feeding on *P. aeruginosa* PAO1 or *S. enterica* MST1.

**Table S3**. Pathogen specific and shared differentially expressed genes in each condition.

**Table S4-5**. Small RNAs differentially expressed in pathogenic conditions. Up and downregulated sRNAs in the F1 and F2 of animals feeding on *P. aeruginosa* PAO1 (4) or *S*. *enterica* MST1(5).

**Table S6**. Pathogen specific and shared differentially expressed genes in each condition. (14) upregulated and (15) downregulated genes.

**Table S7**. Putative targets of *mir-243* among genes downregulated in pathogenic conditions.

**Dataset 1.** Differential expression analysis of expressed mRNA genes in two generations of animals exposed to *P. aeruginosa* PAO1 and *S*. *enterica* MST1 by DeSeq and EdgeR.

**Dataset 2.** Differential expression analysis and genomic context of expressed sRNA genes in two generations of animals exposed to *P. aeruginosa* PAO1 and *S*. *enterica* MST1.

## Acknowledgments

We are deeply grateful to Marcia Manterola in the University of Chile, who provided a laboratory in times of need. Without her help the finalization of this work would not have been possible. Ana Maria Pozo facilitated the timely acquisition of key reagents for this work. Some strains were provided by the CGC, which is funded by NIH Office of Research Infrastructure Programs (P40OD010440). This work funded by Millennium Scientific Initiative of the Chilean Ministry of Economy, Development, and Tourism (P029-022-F), Proyecto Apoyo Redes Formacion de Centros (REDES180138), ANID Programa Cooperación Internacional CYTED grant P918PTE 3, CONICYT-USA 0041 and Fondecyt 1131038 to AC. The funders had no role in study design, data collection and interpretation, or the decision to submit the work for publication.

## Author contribution

Conceptualization: MFP and AC

Methodology: ML, MFP, FG and AC

Investigation: CG, ML, MFP, LV, FG and AC

Writing-Original Draft: AC

Writing-Review and Editing: CG, ML, MFP and AC

Funding Acquisition: AC

