## Supplementary material for "Intergenerational pathogen-induced diapause in *C. elegans* is modulated by *mir-243*": Figure S1

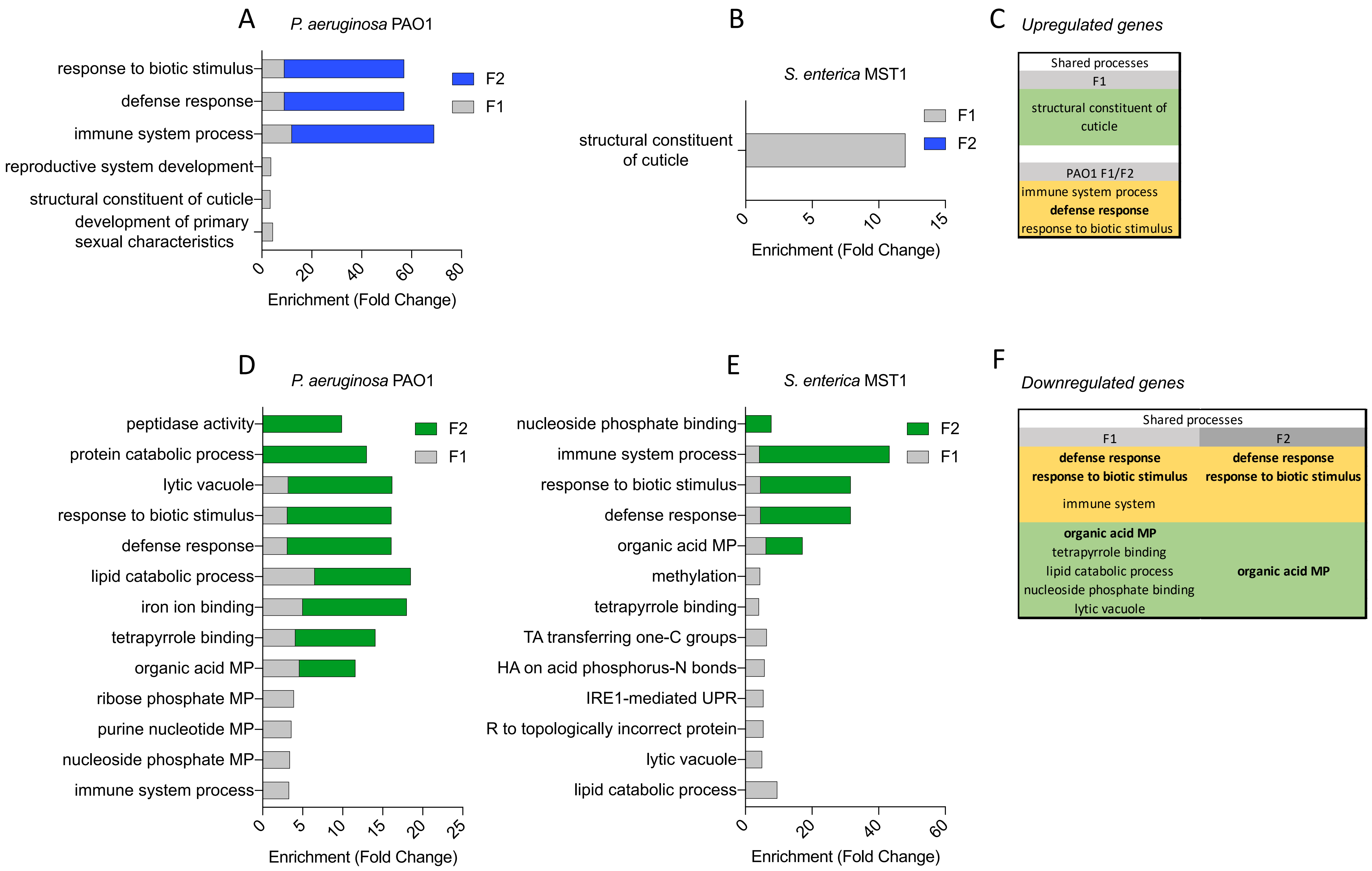


**Fig. S1**. Enrichment by GO term of upregulated (A,B) and downregulated (D,E) in animals feeding on *P. aeruginosa* PAO1 and *S. enterica* serovar Typhimurium MST1 in two generations. C, F Summary of shared GO terms in F1 and F2 in up (C) and downregulated (F) genes. MP, metabolic process; TA, transferase activity; HA, hydrolase activity; N, Nitrogen; C, Carbon; R, response; P, phosphate; UPR, Unfolded Protein Response.
